# Molecular Determinants of Prognosis and Evolution in Diffuse-Lower Grade Astrocytomas

**DOI:** 10.1101/2020.08.02.232918

**Authors:** Mehul Kumar, J Gregory Cairncross, Michael D Blough, Pinaki Bose

**Author notes:** Corresponding Authors: Pinaki Bose, HMRB 354, 3330 Hospital Drive NW, Calgary, Alberta, Canada T2N 4N1, Michael D. Blough, HRIC2A27, 3330 Hospital Drive NW, Calgary, Alberta, Canada T2N 4N1. Funding: The Terry Fox Research Institute and Foundation, the Alberta Cancer Foundation, Genome Canada, Alberta Innovates, and the family of Clark H Smith supported this work.

## Abstract

Low grade astrocytomas (LGAs) are classified based on the mutational status of the isocitrate dehydrogenase (IDH) gene. While IDH wild-type (WT) LGAs evolve rapidly to glioblastoma, mutant tumors generally have a more indolent course. To identify potential drivers of the differential progression of LGAs, we analyzed transcriptomes from The Cancer Genome Atlas. Compared to mutant LGAs, platelet-derived growth factor (PDGF) signaling is enriched in WT cases, and *PDGFA* is the top overexpressed gene in the pathway. Putative mechanisms for differential *PDGFA* expression included copy number gains of chromosome 7 in WT cases and methylation of the *PDGFA* promoter in mutant LGAs. Additionally, we found that high *PDGFA* expression is associated with aneuploidy, immunosuppressive features, and worse prognosis, and that WT LGAs use multiple means to inactivate the p53 pathway to progress to GBM. Our work highlights the contribution of PDGF gene family towards the unique behaviour of LGAs.

**STATEMENT OF SIGNIFICANCE:** This study of gene expression in LGAs suggests that differential regulation of the PDGF pathway may underlie the different natural histories of *IDH* WT and *IDH* mutant LGAs including divergent evolutionary trajectories to GBM. This insight may inspire new therapeutic strategies to suppress the transformation of LGAs to higher-grade cancers.

## INTRODUCTION

Diffuse fibrillary astrocytomas of adults (WHO grades II and III), also referred to as lower grade astrocytomas (LGAs), are a group of deadly brain cancers with unknown etiology. A significant insight into their biology and variable clinical behaviour was gained when isocitrate dehydrogenase (*IDH*) 1 and 2 mutations were discovered in a large proportion of LGAs (1). It soon became apparent that IDH wild-type (WT) LGAs first appeared in older adults and evolved rapidly to glioblastoma (GBM), whereas LGAs with an *IDH* mutation occurred in younger adults, grew slowly, and only sometimes evolved into an end-stage GBM-like cancer. Despite similar histology, *IDH* WT and mutant LGAs had very different natural histories. Soon, additional molecular and biochemical features distinguishing *IDH* WT LGAs from mutant tumors were identified, including amplification of chromosome 7, deletion of chromosome 10, and mutations of the *TERT* gene promoter in *IDH* WT cases, versus loss of the alpha-thalassemia x-linked (*ATRX*) gene, point mutations of the *TP53* gene, and production of the oncometabolite, 2-hydroxyglutarate (2-HG), in *IDH* mutant LGAs (2). But how do these well-characterized molecular alterations mediate their different natural histories?

In an effort to answer this question, we analyzed transcriptome data on all non-1p and 19q co-deleted LGAs in The Cancer Genome Atlas (TCGA). Here, we describe previously unrecognized differences between *IDH* WT and mutant tumors with respect to PDGF signaling, particularly overexpression of *PDGFA* and *PDGFD* driven by promoter methylation and copy number variation (CNV). We also observed that overexpression of *PDGFA*/*PDGFD* is associated with aneuploidy, markers of immunosuppression, and worse outcome. Further, the evolution of WT LGAs to GBM is accompanied by inactivation of multiple components of the p53 pathway. Our data support a role for PDGF genes in the early stages, progression, and prognosis of LGAs.

## RESULTS

### Enrichment of the PDGF pathway and high expression of PDGFA is observed in IDH WT LGAs

To explore putative mechanisms underlying the stark differences in survival of *IDH* WT and mutant LGAs we performed differential expression analysis (DEA) on LGAs stratified by *IDH1*/*2*-mutation status from a filtered TCGA dataset (n = 347). In this assessment we identified 2,175 overexpressed and 517 downregulated genes in WT LGAs versus mutant cases (adjusted *P*-value = 0.001, log_2_(fold change) = 1; Fig. 1A). To assess functional outcomes of differentially expressed genes, we performed canonical Reactome pathway analysis: enriched pathways in WT tumors included ECM deregulation, collagen biosynthesis, and PDGF signaling (Fig. 1B). Although enrichment of ECM pathways was expected (3) given the invasive nature of LGAs, the prominence of the PDGF pathway was of interest because overexpression of *PDGFA* has been implicated in the etiology of *IDH* WT GBM (4) and the PDGFA ligand has been shown to initiate GBM in animal model systems (4,5).

**Figure 1:**
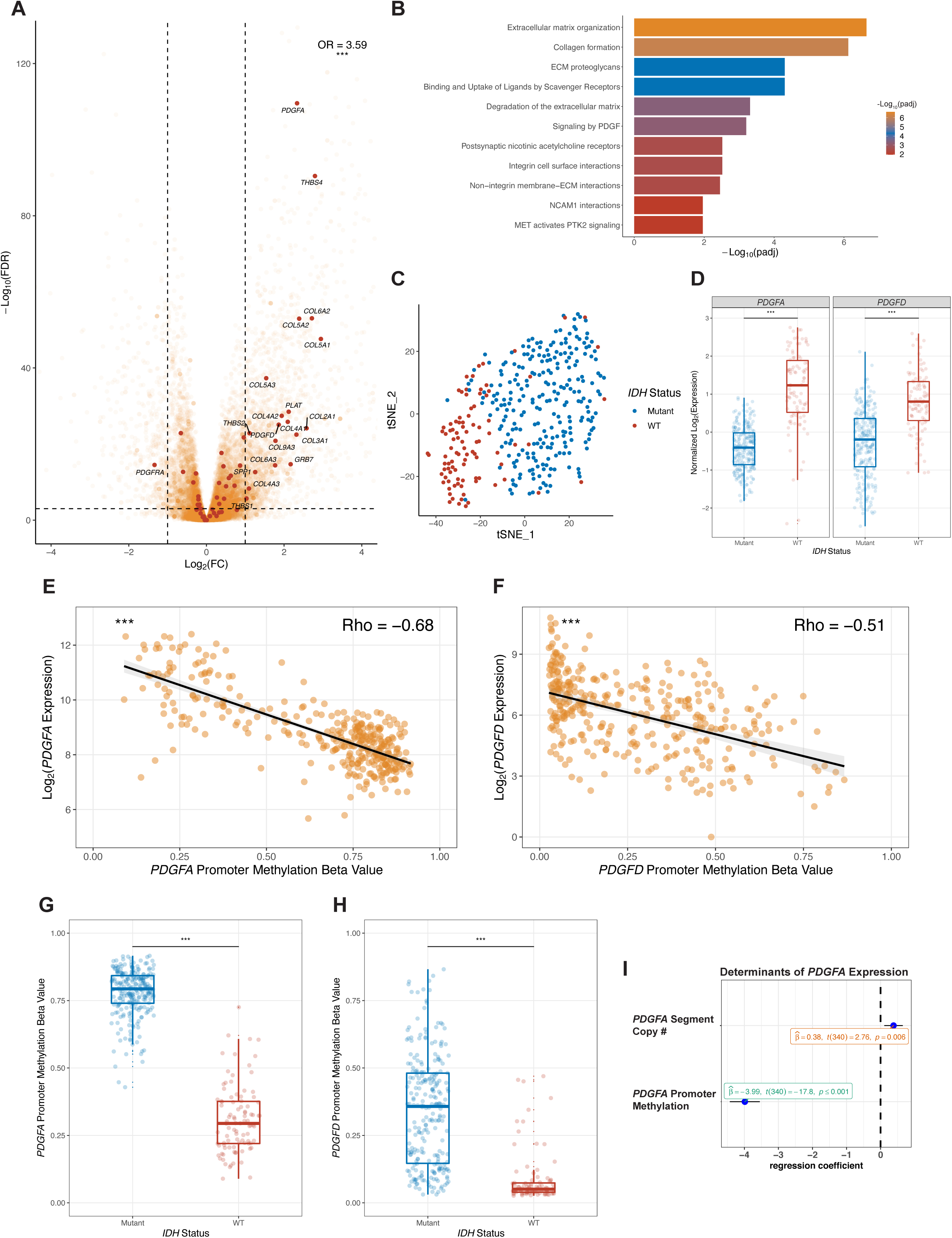
*PDGFA* and *PDGFD* expression is dysregulated in *IDH* WT LGAs. **(A)** Volcano plot showing fold changes for genes differentially expressed between *IDH* WT and *IDH* mutant LGAs. PDGF pathway members are enriched in the overexpressed genes, as shown in maroon. **(B)** Reactome pathway analysis of genes overexpressed in *IDH* WT LGAs reveals the enrichment of ECM-associated genes and the PDGF signaling pathway. **(C)** Unbiased tSNE visualization with gene expression values of PDGF pathway genes suggesting segregation of LGAs by *IDH* mutation status. **(D)** *PDGFA* and *PDGFD* gene expression is significantly elevated in *IDH* WT LGAs, relative to *IDH* mutant LGAs. Scatterplots showing the negative correlation of promoter methylation with **(E)** *PDGFA* and **(F)** *PDGFD* expression across LGAs. Spearman’s Rho values are reported as a measure of effect size. Box plots showing that promoter methylation of **(G)** *PDGFA* and **(H)** *PDGFD* is elevated in *IDH* mutant relative to *IDH* WT LGAs. **(I)** Multivariate linear model showing the independent association of *PDGFA* expression with *PDGFA* promoter methylation and copy number of the segment containing *PDGFA* on chromosome 7. OR – Odds Ratio. (*** *P* < 0.001).

The most differentially expressed gene in the family was *PDGFA* (Fig. 1A and C). *PDGFA*, like *PDGFD*, was significantly upregulated in *IDH* WT LGAs compared to mutant LGAs (Fig. 1A and C), while its receptor, *PDGFRA*, was significantly overexpressed in mutant cases (Fig. 1A). *PDGFB, PDGFC, and PDGFRB* were not differentially expressed, but the pattern of expression of the PDGF gene family could differentiate WT from mutant LGAs in an unsupervised analysis using t-stochastic neighbor embedding (tSNE; Fig. 1D) or hierarchical clustering (Fig. S1).

We then explored mechanisms underlying the differential expression of *PDGFA* and *PDGFD* in WT and mutant LGAs. Aware that hypermethylation is a feature of *IDH* mutant LGAs (6), we first assessed whether promoter methylation was associated with *PDGFA/PDGFD* expression, and documented a strong negative correlation between expression and methylation for both genes across all LGAs (Fig. 1E and F); *PDGFA* and *PDGFD* promoter methylation were absent in *IDH* WT LGAs, but present in mutant cases (Fig. 1G and H, respectively). The negative correlation between expression and methylation was preserved when WT and mutant LGAs were analyzed separately (Fig. S2A-D), indicating for the first time that promoter methylation may be an important regulatory mechanism of *PDGFA*/*PDGFD* expression in both LGA subtypes.

We next assessed the correlation between gene expression and chromosome copy number to determine if gains of chromosome 7 (*PDGFA*) and chromosome 11 (*PDGFD*) were associated with differential gene expression. As reported by others (7), we found that a significantly higher proportion of *IDH* WT LGAs displayed amplification of the portion of chromosome 7 containing *PDGFA* compared to mutant LGAs (Fig. S3A). *PDGFD* segment amplification is not a prominent feature in LGAs (Fig. S3B). Furthermore, the absolute copy number of the *PDGFA* containing segment on chromosome 7 significantly correlated with *PDGFA* expression in *IDH* WT LGAs (Fig. S3C), but not mutant cases (Fig. S3D). In multivariate linear regression analysis, both *PDGFA* promoter methylation and copy number of the *PDGFA* containing segment were associated with *PDGFA* expression (Fig. 1I).

Together, these results reveal a previously unidentified mechanism by which PDGF signaling may be regulated in LGAs. In *IDH* WT LGAs, absence of promoter methylation of *PDGFA* and *PDGFD* and amplification of chromosome 7 contribute to higher gene expression. Whereas in *IDH* mutant LGAs, hypermethylation of the *PDGFA* and *PDGFD* promoters and absence of chromosome 7 amplification lead to the decreased expression of *PDGFA* and *PDGFD*.

### Gene expression, promoter methylation of PDGFA/PDGFD and amplification of PDGFA are significantly associated with prognosis LGA

Survival analyses and Kaplan-Meier (KM) curves were used to assess whether gene expression and/or promoter methylation of *PDGFA* and PDGFD were prognostic factors for LGAs. High PDGFA expression was associated with shorter overall survival (OS), disease specific survival (DSS) and progression-free interval (PFI) (Fig. 2A-C). These results were confirmed in two additional datasets (GSE16011; Fig. 2D, <u>Re</u>pository for <u>M</u>olecular <u>BRA</u>in <u>N</u>eoplasia <u>D</u>a<u>T</u>a (REMBRANDT Project; Fig. 2E) supporting the validity of *PDGFA* expression as a prognostic biomarker in LGA. Similar associations were observed for *PDGFD* expression (Fig. S4A-E). Lower *PDGFA/PDGFD* promoter methylation (Fig. S5A-F) and amplification of the chromosome segment containing *PDGFA* (Fig. 2F-H) were also associated with shorter OS, DSS and PFI. All survival associations were initially confirmed to be significant in univariate Cox proportional hazards models with the continuous variables as covariates (data not shown). These data suggest that mechanisms regulating *PDGFA/PDGFD* expression may impact the biology and clinical behavior of LGA.

**Figure 2:**
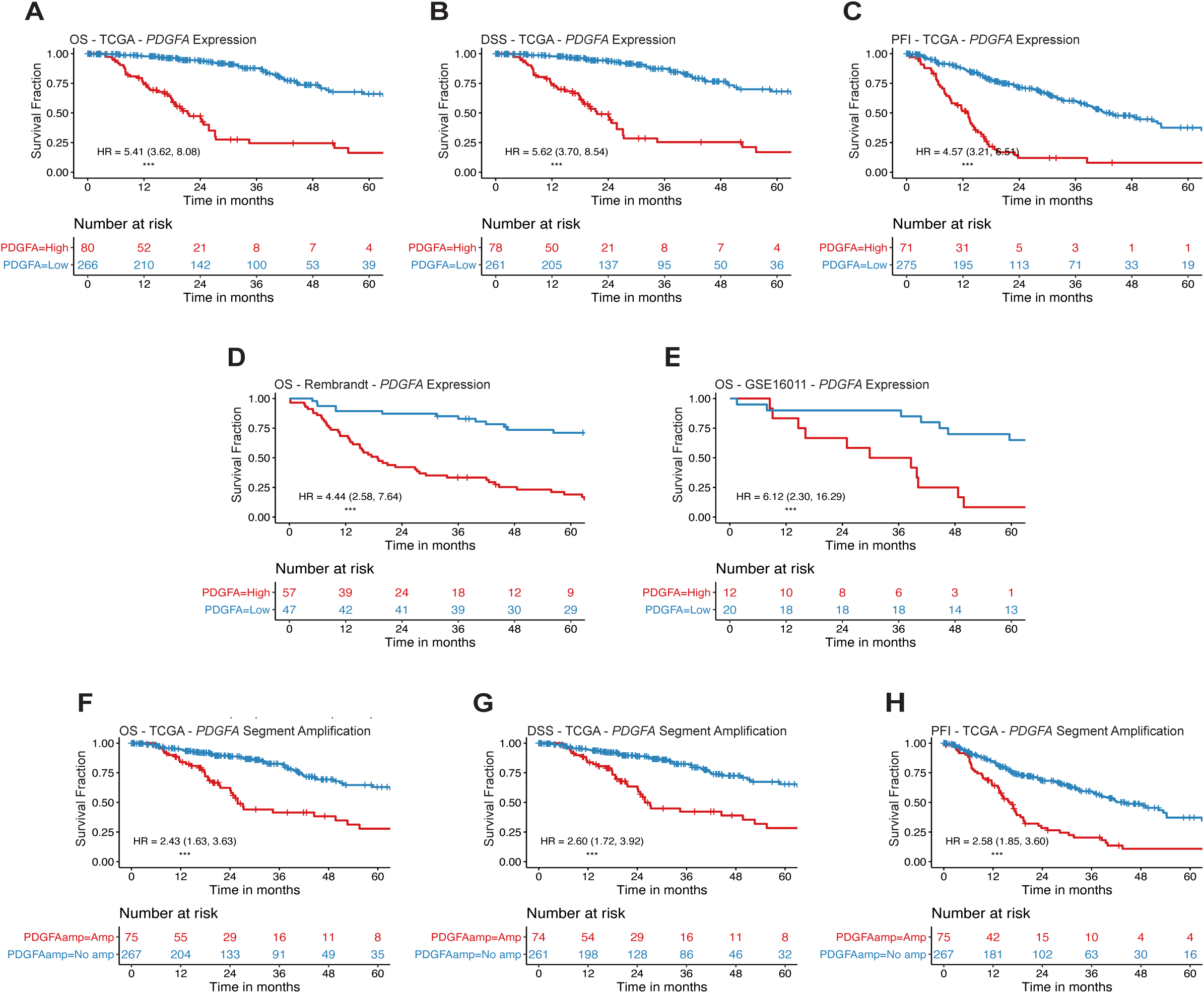
*PDGFA* expression and amplification status are associated with worse prognosis in LGAs. KM survival curves for **(A)** OS **(B)** DSS and **(C)** PFI of TCGA LGA patients divided into low and high-risk groups based on *PDGFA* expression. KM survival curves validating the association between *PDGFA* expression of the tumor and overall survival of the patient in LGA samples from **(D)** the REMBRANDT and **(E)** GSE16011 datasets. KM survival curves for **(F)** OS **(G)** DSS and **(H)** PFI from TCGA LGA patients divided into risk groups based on whether the chromosomal segment containing the *PDGFA* locus is amplified or not-amplified. Hazard ratios (HR) and their respective 95% confidence intervals from univariate Cox proportional hazard analysis of the dichotomized expression groups are reported for each KM curve. (*** *P* < 0.001).

### PDGFA and PDGFD gene expression and IDH WT status are associated with aneuploidy and markers of immunosuppression

Given the poor prognosis of *IDH* WT and *PDGFA* overexpressing LGAs, we then assessed additional biological features that might explain their propensity for GBM-like behavior. Having recently reported that *in vitro* exposure to PDGFA leads to chromosomal instability in neural progenitor cells (5) we assessed aneuploidy in LGAs in relation to *IDH* mutational status and *PDGFA* expression. In this analysis we observed that *IDH* WT LGAs were significantly more aneuploid than their *IDH* mutant counterparts (Fig. 3A). Further, survival analyses revealed that a high aneuploidy score (AS) was associated with worse OS, DSS, and PFI (Fig S6A-C). In the multivariate analysis, both AS and *IDH* status remained independent predictors of survival in LGA (Fig. 3B). We further observed that aneuploidy was a distinguishing feature of LGAs that expressed high levels of *PDGFA* and *PDGFD*; both *PDGFA* and *PDGFD* expression were significantly associated with AS (Fig. 3C and D). These analyses indicate that AS has prognostic value independent of *IDH* status in LGAs, and that aneuploidy is associated with high expression of *PDGFA* and *PDGFD* genes.

**Figure 3.**
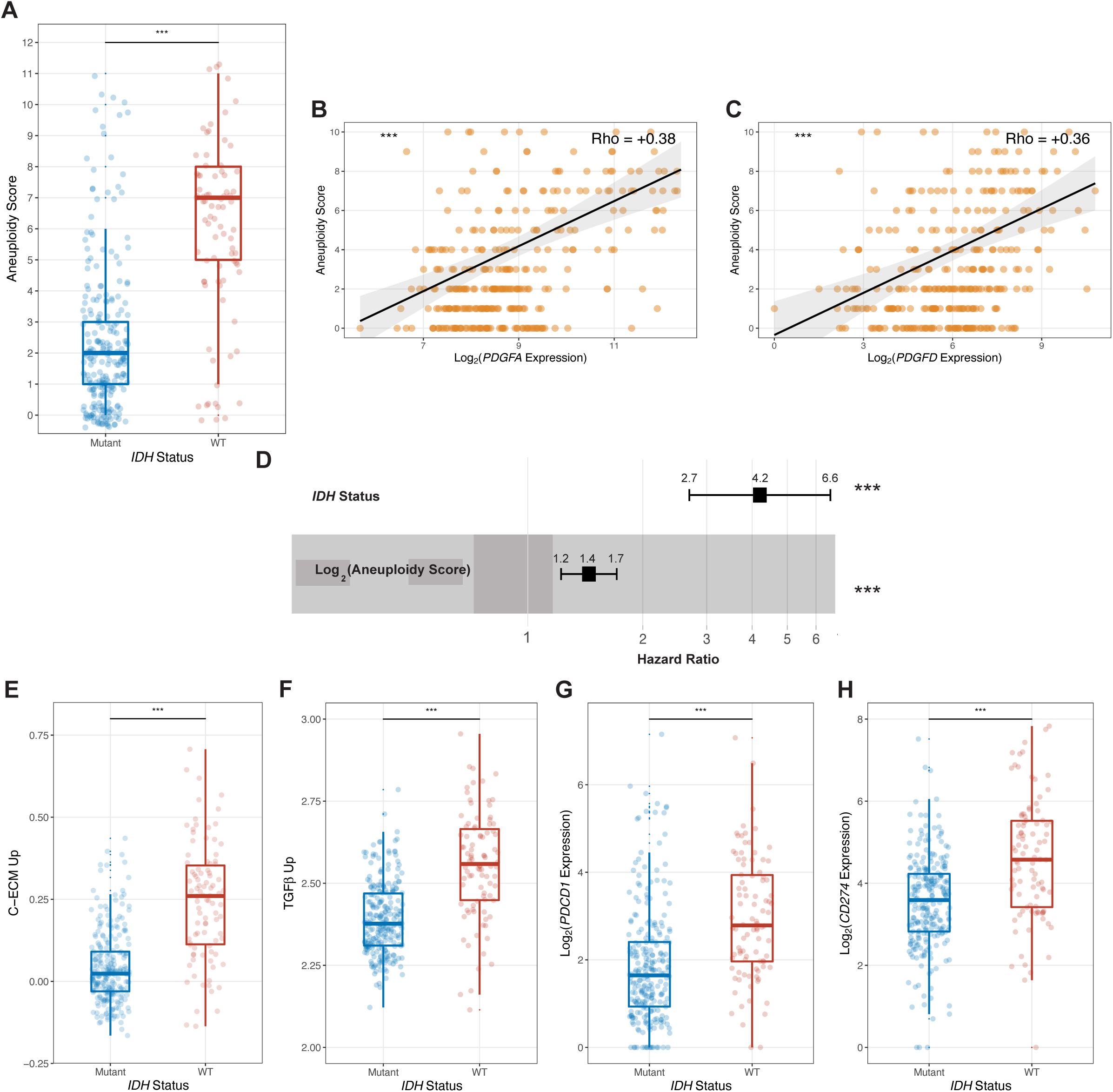
*PDGFA* and *PDGFD* expression are associated with markers of immuno-suppression in LGA patients. **(A)** Box plots depicting the quantification of aneuploidy scores (AS) in *IDH* WT and mutant LGAs. Scatterplots showing the correlations between AS and **(B)** *PDGFA* expression, **(C)** *PDGFD* expression; Spearman’s Rho is reported. **(D)** Multivariate hazard ratio (HR, with 95% CI) derived from a Cox proportional hazards regres-sion model for OS using *IDH* mutation status and Log2(AS+1) as covariates; HR > 1 indicates that *IDH* WT status and high AS are associated with worse OS. Box plots depicting the quantification of **(E)** C-ECM up genes, **(F)** TGF-β upregulated target genes, **(G)** *PDCD1* (PD-1) and **(H)** *CD274* (PD-L1) in *IDH* WT and mutant LGAs. (*** *P* < 0.001).

We then assessed LGA capacity for immune evasion, a hallmark of cancer associated with poor prognosis (8,9). As observed in the pathway enrichment analysis, ECM genes were among those upregulated in WT LGAs (Fig. 1B). This is a compelling observation considering that we have recently reported that ECM dysregulation is an effector of TGF-β-induced immunosuppression in the tumor microenvironment (10). To explore further, we investigated immune suppression in LGAs with respect to their *IDH* mutational status, and documented that WT LGAs had significantly higher expression of TGF-β upregulated target genes (Fig. 3E) and cancer-associated ECM (C-ECM) genes (Fig. 3F). The expression of immunosuppressive checkpoint genes such as *PDCD1* (encodes PD-1) and *CD274* (encodes PD-L1) were also increased in *IDH* WT LGAs (Fig. 3G and H, respectively). These data suggest *IDH* WT LGAs have the capacity to suppress the local immune response.

### Evolution of IDH WT LGAs involves sustained PDGFA overexpression and progressive inactivation of the p53 pathway

To further explore the significantly more aggressive behavior of WT LGAs and to better understand how they evolve to higher-grade cancers, we evaluated the expression of *PDGFA* and *PDGFD* genes in low (WHO grade II), intermediate (grade III), and high-grade astrocytomas (GBM; grade IV) and found that high *PDGFA*/*PDGFD* expression was a constant feature of the *IDH* WT disease, irrespective of histological grade (Fig 4A and B). These observations reveal that high *PDGFA* expression is an early feature of *IDH* WT LGAs that persists as these cancers evolve from grade 2 lesions, to grade 3 lesions, and on to GBM.

**Figure 4:**
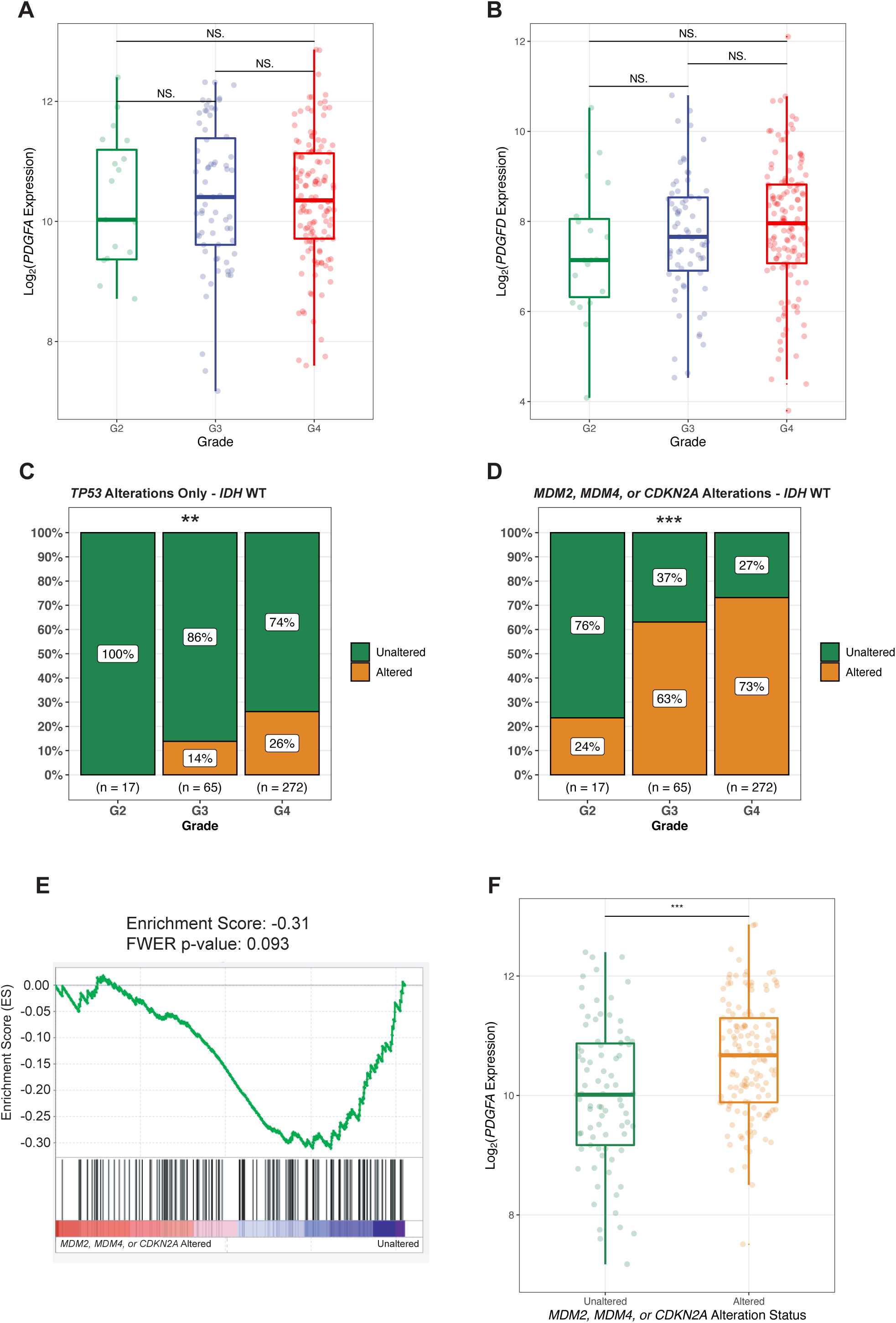
The evolution of *PDGFA*/*PDGFD* overexpressing *IDH* WT LGAs to higher-grade disease is accompanied by a progressive increase in TP53 pathway mutations. Box plots depicting the quantification of **(A)** *PDGFA* and **(B)** *PDGFD* expression across grade in *IDH* WT LGAs. Bar plots showing the proportion of *IDH* WT LGAs harbouring alterations by grade in **(C)** the *TP53* gene and **(D)** either of these genes: *MDM2, MDM4* or *CDKN2A*. **(E)** Gene set enrichment analysis plot, enrichment score (ES) and family wise error rate (FWER) p-value showing the depletion of a p53 target gene set in *MDM2*/*MDM4*/*CDKN2A* altered *IDH* WT LGA patients. **(F)** Box plots depicting the quantification of *PDGFA* expression in *MDM2*/*MDM4*/*CDKN2A* altered versus unaltered *IDH* WT LGA patients. (*** *P* < 0.001; NS: not significant, *P* > 0.1).

We then assessed the mutational status of the p53 pathway, because loss or inactivation of *TP53* has been hypothesized to cooperate with PDGF signaling to promote *IDH* WT GBM (4), and because *TP53* compromise (i.e., null or heterozygous) is a prerequisite for PDGFA-mediated transformation of neural progenitor cells (5). We first assessed single nucleotide variants (SNVs) in *TP53*. Unlike *IDH* mutant LGAs (Fig. S8A), which had a very high rate of SNVs at all WHO grades, we observed no *TP53* SNVs in grade II *IDH* WT LGAs. *TP53* SNVs were only seen in grade 3 *IDH* WT LGAs and *IDH* WT GBMs (Fig. 4C).

Aware that the *TP53* pathway can be inactivated by mechanisms other than gene mutation, we assessed copy number variants (CNVs) of *CDKN2A*, which encodes the p53-positive regulator p14ARF, and variants of MDM2 and MDM4, p53-negative regulators (11). Here, we observed a progressive increase in the frequency of CDKN2A and MDM2/MDM4 alterations with increasing grade in WT LGAs (Fig. 4D). This analysis implies that p53 pathway disruption accompanies progression to GBM in *IDH* WT LGAs in virtually all cases (Fig. S8C).

We then sought to confirm whether deletion of *CDKN2A* and amplification of *MDM2/MDM4* deregulated the p53 pathway in *IDH* WT LGAs. Gene set enrichment analysis (GSEA) using a list of *TP53* target genes confirmed a trend towards significance (*P* = 0.09) for the depletion of this gene set in *CDKN2A/MDM2*/*MDM4* altered WT LGAs (Fig 4E). Interestingly, WT LGAs with alterations in *CDKN2A/MDM2*/*MDM4* showed increased expression of *PDGFA* versus unaltered tumors (Fig. 4F). These data indicate that the first step in the progression of WT grade 2 LGA, to grade 3 LGA, and beyond to GBM, involves inactivation of the p53 pathway by one of several mechanisms, often an early alteration in either *CDKN2A or MDM2*/*MDM4* on a background of high *PDGFA* expression.

## DISCUSSION

The prognostic differences between *IDH* WT and *IDH* mutant LGAs are well known, but the biology that lies beneath these differences, and the specific role of *IDH* status in orchestrating their dissimilar behaviors is still unclear. Our analyses raise the intriguing possibility that *IDH* status acts through differential expression of PDGF pathway genes, especially PDGFA, to create such differences. Our data also raises the further possibility that these differences are related to patterns of methylation of PDGF pathway components. Our work reveals for the first time that PDGF pathway genes are markedly enriched within the pool of differentially expressed genes in WT versus mutant LGAs. We have found that *PDGFA* and *PDGFD* are overexpressed in *IDH* WT LGAs, while their promoter regions are methylated in mutant tumors. Amplification of chromosome 7, which is only seen in *IDH* WT cases and includes the *PDGFA* locus, is a second important difference between WT and mutant LGAs, and emerges as is a putative mechanism for the differential regulation and overexpression of *PDGFA* in WT cases. Furthermore, *PDGFA* expression, promoter methylation of *PDGFA/PDGFD*, and amplification of chromosome 7 are associated with poor survival. Consistent with worse outcomes, *PDGFA* and *PDGFD* expression are correlated with high aneuploidy scores and markers of immune evasion in *IDH* WT LGAs.

A further difference between *IDH* WT and *IDH* mutant LGAs has emerged from this study, and speaks to their very different biological characteristics. This difference pertains to the timing and nature of the p53 pathway alterations found in WT versus mutant LGAs. *TP53* point mutations (i.e., SNVs), primarily located in the DNA binding domain, were an early, frequent, and constant feature of *IDH* mutant LGAs, whereas there were no point mutations of the *TP53* gene in WHO grade 2 *IDH* WT LGAs. Point mutations in WT LGAs were seen at later stages of the disease in grade 3 and 4 (i.e., GBM) cases only. Instead of point mutations, the p53 pathway was more often inactivated by other mechanisms in WT LGAs, primarily by deletions of *CDKN2A* and overexpression of *MDM2* and *MDM4*. Indeed, deletions of *CDKN2A* were the earliest alteration affecting the p53 pathway in *IDH* WT cases. Such differences have not been fully appreciated.

In context of the current literature, these data emphasize the scope of genomic reprogramming that occurs in the diffuse astrocytic gliomas of adults related to the presence or absence of an *IDH* mutation, and points to the potentially important roles of the PDGF and p53 pathways in the behavior of LGAs. Such information might be incorporated into future therapeutic strategies or risk assessments. For example, demethylating agents have entered clinical trials for *IDH* mutant LGAs, yet published data pointing to *PDGFA* overexpression as a potential initiator of *IDH* WT GBM (4) raises the concern that its silencing by promoter methylation in *IDH* mutant cases might be naturally protective against GBM. Perhaps caution should be exercised when using such agents as therapies for mutant LGAs, lest the restoration of *PDGFA* expression inadvertently hastened the evolution of *IDH-TP53* mutant LGAs to GBM (5).

Our analysis also reveals that another member of the PDGF family, *PDGFD*, is significantly upregulated in *IDH* WT LGAs. Although there is very little evidence implicating *PDGFD* in the pathogenesis of LGA, heterodimers and homodimers of *PDGFD* bind to PDGFRα and PDGFRβ, respectively (12,13). *PDGFRA* (the gene encoding PDGFRα) is expressed at lower levels in WT LGAs, while *PDGFRB* (the gene encoding PDGFRβ) is upregulated in *IDH* WT cases. Perhaps *IDH* WT LGAs have developed a mechanism that allows for the modulation and activation of both PDGF receptors. Speculation aside, our observations suggest that *PDGFA*, and other PDGF family members, are potential mediators of the different clinical behaviors of IDH *WT* and mutant LGAs, and that WT and mutant tumors trace distinct evolutionary trajectories to GBM. Whether PDGF inhibitors might be effective therapies for *IDH* WT LGAs is unknown and currently very difficult to test. Testing will require new tools for early detection of WT cases, because in a new *in vitro* model of GBM initiation by *PDGFA*, pathway inhibitors are effective only at the earliest stages of transformation (5).

## METHODS

### Data Analysis and Statistical Tests

Data wrangling, cleaning, and analyses were performed on R version 4.0.0. All statistical tests were two-sided.

### Datasets Used

TCGA clinical data for LGG and GBM cases was downloaded from supplementary table 1 in Ceccarelli *et al*. (14). This dataset was utilized for annotated information on the grade, *IDH* status and 1p/19q co-deletion status of TCGA gliomas. Pan-cancer (including TCGA-GBM/LGG) survival data was downloaded from supplementary table 1 by Liu *et al*. (15). *TP53* pathway genes (*TP53, MDM2, MDM4, or CDKN2A*) were queried for mutations and copy number alterations in the TCGA-GBM/LGG dataset on the cBio Cancer Genomics Portal (cbioportal.org) (16). The corresponding raw dataset for the OncoPrint generated by cBioPortal was downloaded for analysis on a third-party platform. AS for TCGA-GBM/LGG samples were acquired from supplementary table 2 provided alongside the study by Taylor *et al*. (17).

### Gene Expression, Copy Number, and Methylation Datasets

Normalized level 3 RSEM RNA-seq data, segmented copy number data from SNP6 arrays, and Infinium 450k methylation array data for TCGA-GBMLGG samples was downloaded from the Broad GDAC Firehose (https://gdac.broadinstitute.org). For copy number analysis, probes were filtered to those overlapping the region containing the *PDGFA* (chromosome 7 between 536897 bp and 559481 bp) or *PDGFD* (chromosome 11 between 103777914 bp and 104035027 bp) genes. Absolute copy number values were computed by transforming segment means (absolute copy # = 2 ×2^(segment mean)^). Methylation probes cg15454385 and cg03145963 were used as the representative probes to study *PDGFA* and *PDGFD* promoter methylation status, respectively.

### Filtering of Gliomas, and Classification of LGA

Depending on the analysis requirements, filters were utilized to select glioma samples based on their grade or molecular alteration status (*IDH* status, 1p/19q codeletions, or *TP53* pathway alterations). LGAs represent WHO grade II/III gliomas without 1p/19 codeletions.

### Differential Expression Analysis of IDH WT LGAs vs IDH Mutant LGAs in TCGA

Level 3 RNA-Seq by Expectation-Maximization (RSEM) data was downloaded for TCGA-GBM/LGG samples from GDAC Firehose (https://gdac.broadinstitute.org). Samples were filtered for tumors and subsequently for LGAs, as per the classification described above. Samples classified as NA for *IDH* status in the clinical dataset were omitted from the analysis. Differential expression analysis was performed between *IDH* WT and IDH mutant LGAs using the DESeq2 package (18). A differentially expressed gene (DEG) list was generated with an adjusted *P*-value threshold of 0.001 and log_2_(fold change) threshold of +1.

### Pathway Enrichment Analysis of DEGs between IDH WT and IDH Mutant LGAs

Pathway enrichment analysis was performed on a smaller DEG list (with an adjusted *P*-value threshold of 0.001 and log2(fold change) threshold of +2) using the ReactomePA package (19). As multiple pathway hits were related to collagen, scavenger receptors, and acetylcholine receptors, these pathway hits were collapsed into one pathway hit each.

### Hierarchical Clustering and t-Stochastic Neighbor Embedding (tSNE)

RSEM normalized gene expression for PDGF family genes (PDGFA, PDGFB, PDGFC, PDGFD, PDGFRA, PDGFRB) were log_2_ transformed (log_2_(RSEM + 1)) and subsequently converted to Z-scores. LGA samples were hierarchically clustered based on Euclidean distances. Heatmap construction and hierarchical clustering was performed using the pheatmap R package. Exact 2-D tSNE dimensionality reduction was performed on the z-scores of the log-transformed PDGF family gene expression values from LGAs. A principle component analysis (PCA) step with 100 dimensions was performed prior to tSNE. Perplexity for the tSNE was set to 20. tSNE was performed using the Rtsne R package.

### Gene Set Enrichment Analyses

*TP53* pathway genes were identified from the Molecular Signature Database (MSigDB v6.2, C2 collection; https://www.gsea-msigdb.org/gsea/msigdb/genesets.jsp?collection=C2). The list of TGF-β upregulated target genes and cancer-associated ECM (C-ECM) genes were downloaded from the supplementary material in the study by Chakravarthy *et al*. (10). These gene sets were used to compute single sample gene set enrichment analysis (ssGSEA) scores using the gene set variation analysis (GSVA) R package (20,21). For a pre-defined set of genes, ssGSEA calculates an enrichment score based on coordinately enriched and depleted gene expression for each case. Gene set enrichment analysis (GSEA) was performed with the Broad GSEA 4.0.1 software. GSEA permutation type was set to “phenotype” and 1000 permutations were performed.

### Survival Analyses and Kaplan-Meier Visualizations

Cox-proportional hazards models were fit on R with the survival package. Differences in surviving fractions between groups were visualized via Kaplan-Meier curves generated using the survminer R package. Cut-points for continuous variables were identified by the method noted by Contal and O’Quigley (22).

### Validation Cohorts

Gene expression and clinical data from two additional glioma datasets were obtained (GSE16011 and REMBRANDT). For both datasets, gliomas were filtered to contain only LGAs (i.e., WHO Grade II or III).

### Statistical Visualizations

Exploratory graphs with statistical information and bar-plots were generated using the ggstatsplot R package.

## AUTHORS CONTRIBUTIONS

Conception and design: M. Blough, J.G. Cairncross and P Bose

Development of methodology: M. Kumar, M. Blough and P. Bose

Analysis and interpretation of data (e.g., statistical analysis, biostatistics, computational analysis): M. Kumar and P. Bose

Writing, review, and/or revision of the manuscript: All Authors

Study supervision: M. Blough and P. Bose

---

**Figure S1:**
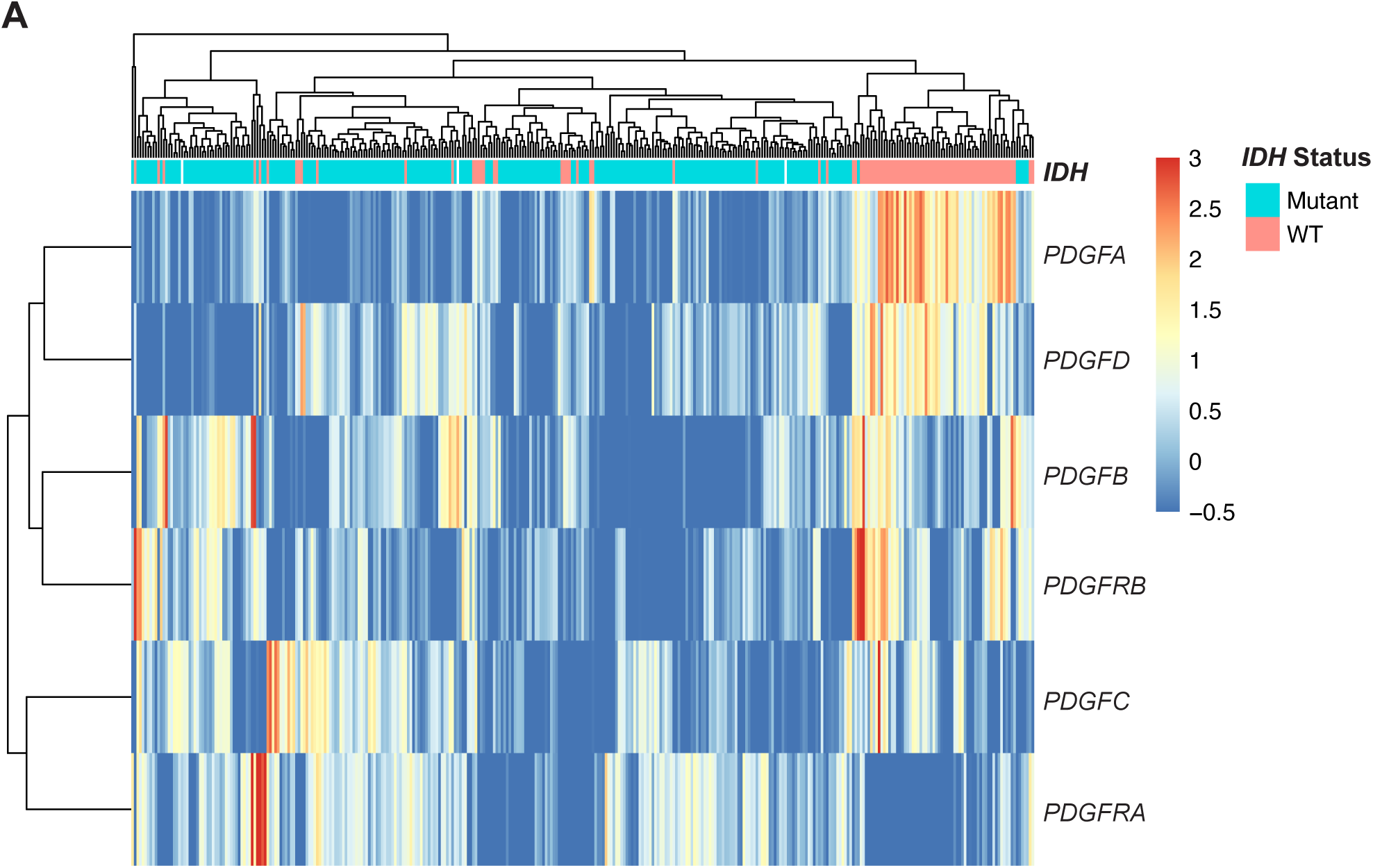
Unsupervised hierarchical clustering of *IDH* WT and mutant LGA samples based on the expression of PDGF family genes. Values are z-scores of log2 transformed RSEM normalized counts. Colours represent relative expression levels of a gene within each row; red represents gene upregulation and blue represents gene downregulation

**Figure S2:**
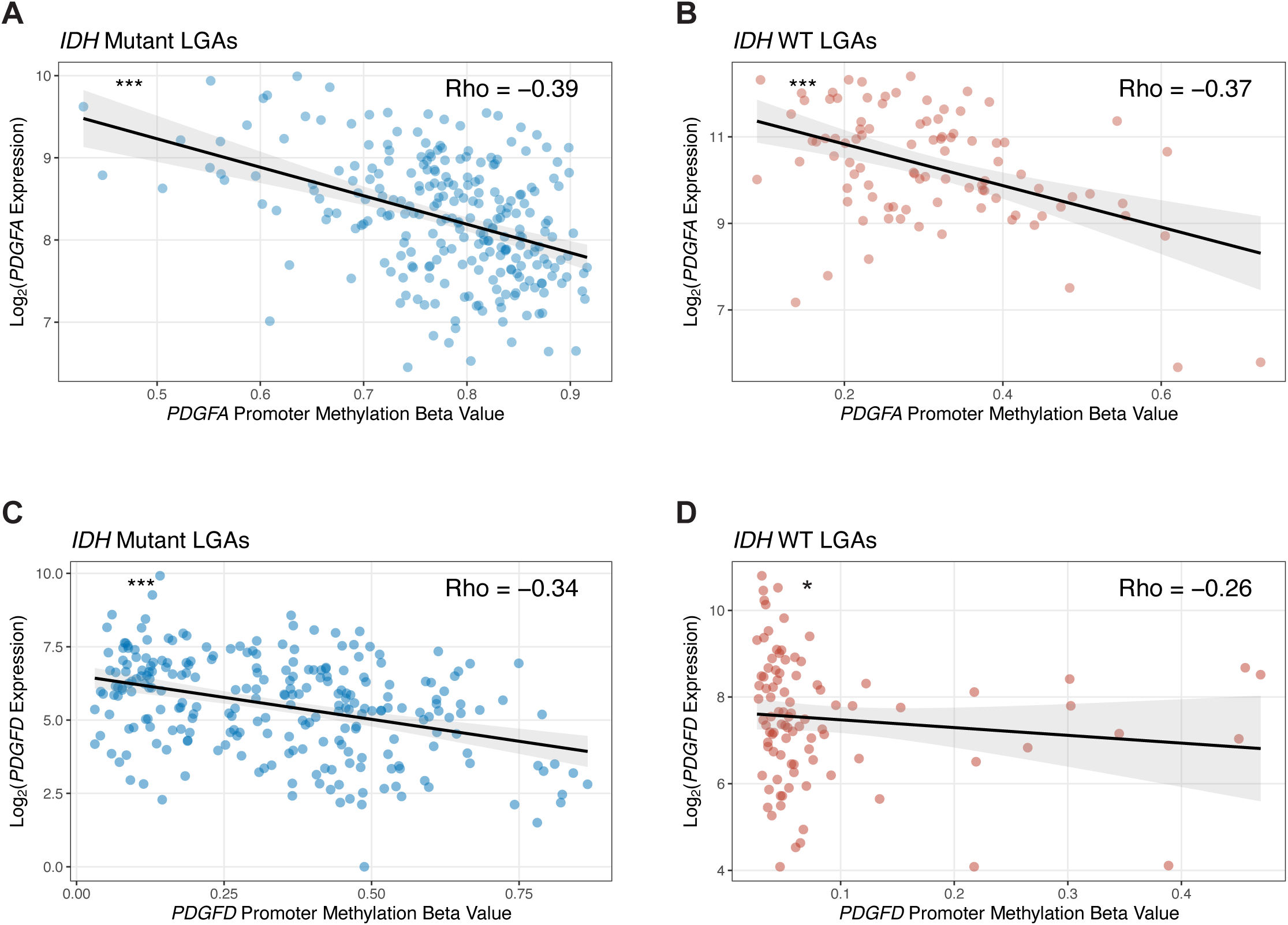
*PDGFA* and *PDGFD* expression is negatively regulated by promoter methylation. Scatterplots showing the negative correlation between *PDGFA* promoter methylation and *PDGFA* expression in **(A)** *IDH* mutant and **(B)** *IDH* WT LGAs. Scatterplots showing the negative correlation between *PDGFD* promoter methylation and *PDGFD* expression in **(C)** *IDH* mutant and **(D)** *IDH* WT LGAs. Spearman’s Rho values are reported. (* *P* < 0.05, *** *P* < 0.001). expression groups are reported for each KM curve. (*** *P* < 0.001).

**Figure S3:**
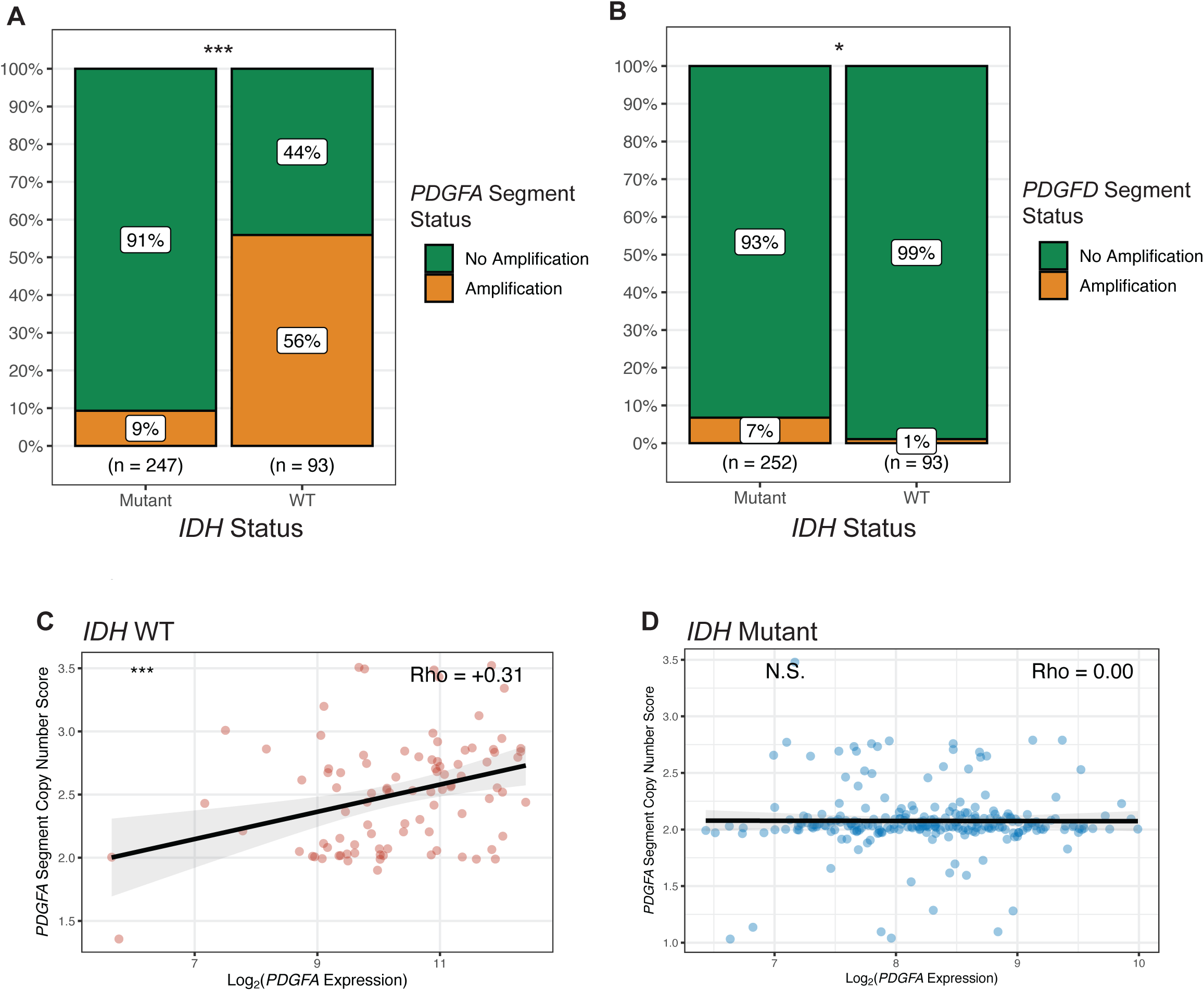
*PDGFA* but not *PDGFD* expression is regulated by chromosomal amplification. Bar plots showing the proportions of samples with **(A)** *PDGFA* and **(B)** *PDGFD* segment amplification in *IDH* mutant and *IDH* WT LGAs. Scatterplots showing the correlation between absolute copy number of the chromosomal segment containing *PDGFA* and *PDGFA* expression in **(C)** *IDH* WT and **(D)** *IDH* mutant LGAs. (* *P* < 0.05, *** *P* < 0.001; N.S.: not significant, *P* > 0.1).

**Figure S4:**
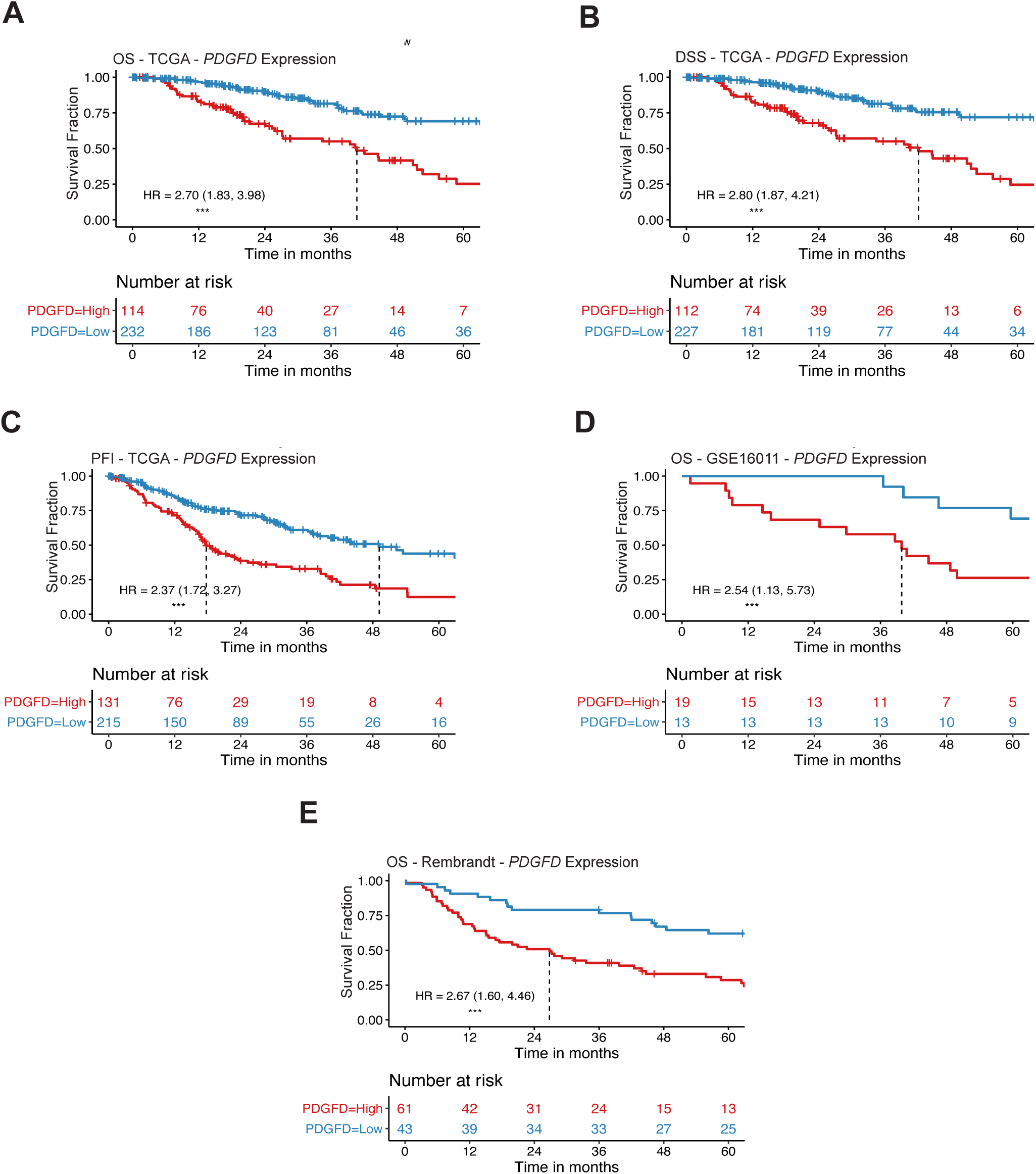
*PDGFD* expression is associated with significantly worse prognosis in LGA. KM survival curves for **(A)** OS **(B)** DSS and **(C)** PFI from TCGA LGA patients divided into low and high-risk groups based on *PDGFD* expression. KM survival curves validating the association between *PDGFD* expression and OS in LGA samples from **(D)** GSE16011 and **(E)** the REMBRANDT datasets. Hazard ratios (HR) and their respective 95% confidence intervals from univariate Cox proportional hazard analysis of the dichotomized expression groups are reported for each KM curve. (* *P* < 0.05, *** *P* < 0.001).

**Figure S5:**
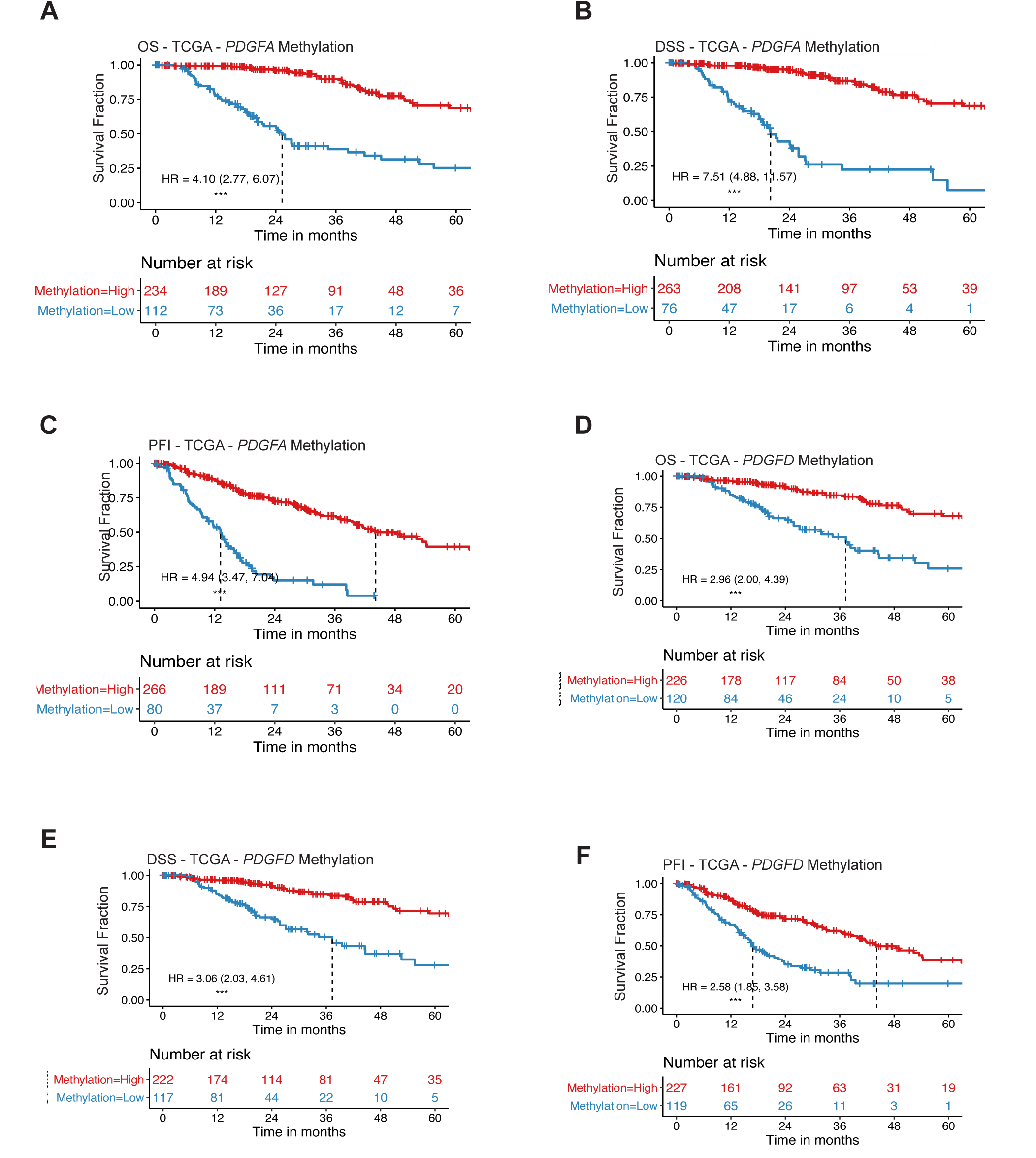
Increased *PDGFA* and *PDGFD* promoter methylation is associated with improved prognosis in LGA. KM survival curves for **(A)** OS **(B)** DSS and **(C)** PFI from TCGA LGA patients divided into low and high-risk groups based on *PDGFA* promoter methylation. KM survival curves for **(D)** OS **(E)** DSS and **(F)** PFI from TCGA LGA patients divided into low and high-risk groups based on *PDGFD* promoter methylation. Hazard ratios (HR) and their respective 95% confidence intervals from univariate Cox proportional hazard analysis of the dichotomized expression groups are reported for each KM curve. (*** *P* < 0.001).

**Figure S6:**
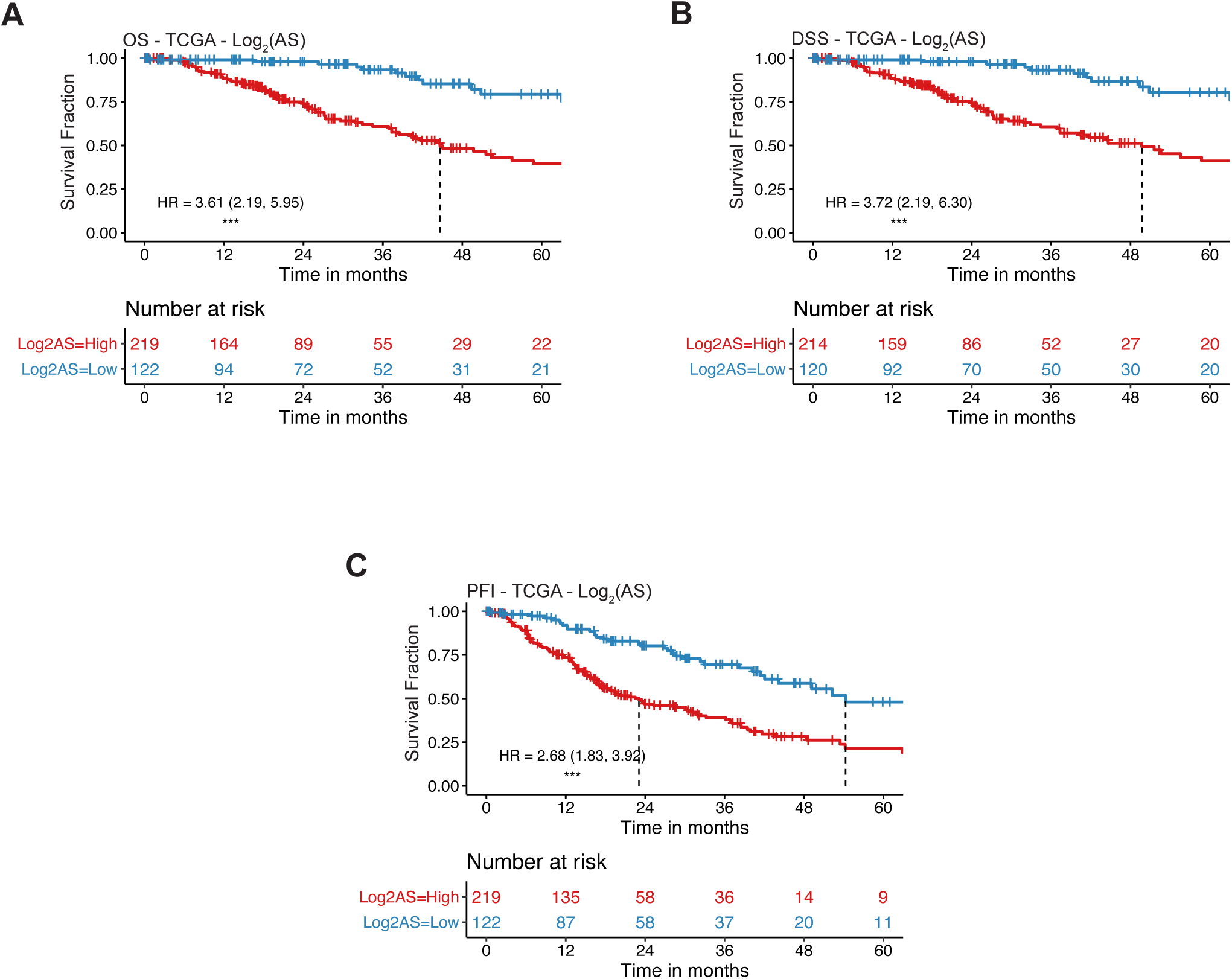
Increased aneuploidy is associated with significantly worse prognosis in LGA. KM survival curves for **(A)** OS **(B)** DSS and **(C)** PFI from TCGA LGA patients divided into low and high-risk groups based on AS. Hazard ratios (HR) and their respective 95% confidence intervals from univariate Cox proportional hazard analysis of the dichotomized expression groups are reported for each KM curve. (*** *P* < 0.001).

**Figure S7:**
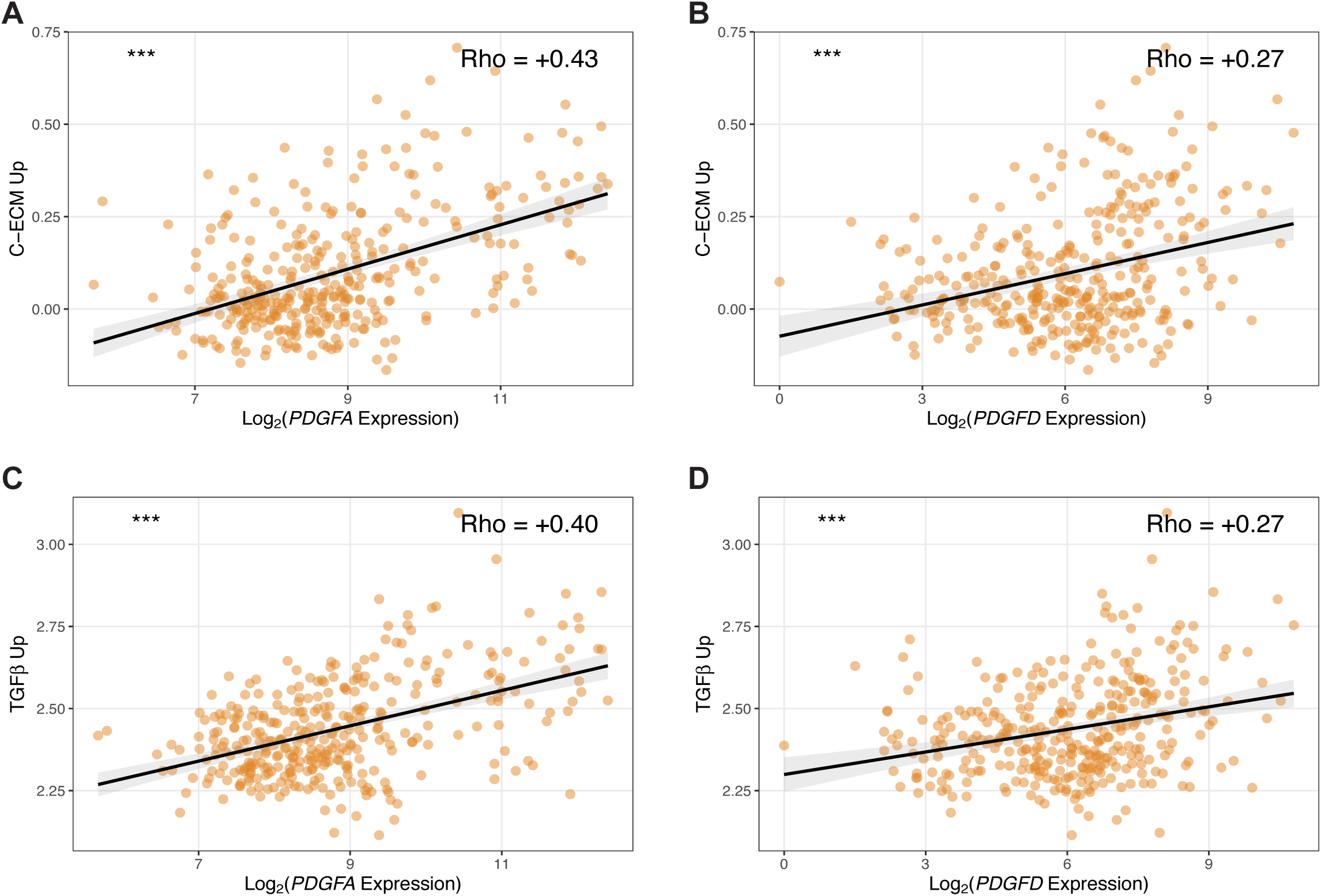
*PDGFA* and *PDGFD* expression is associated with markers of immunosuppression. Scatterplots showing the positive correlation of C-ECM upregulated genes with **(A)** *PDGFA* and **(B)** *PDGFD* expression. Scatterplots showing the positive correlation of TGF-βupregulated target genes with **(C)** *PDGFA* and **(D)** *PDGFD* expression. Spearman’s Rho values are reported. (* *P* < 0.05, *** *P* < 0.001; N.S.: not significant, *P* > 0.1).

**Figure S8:**
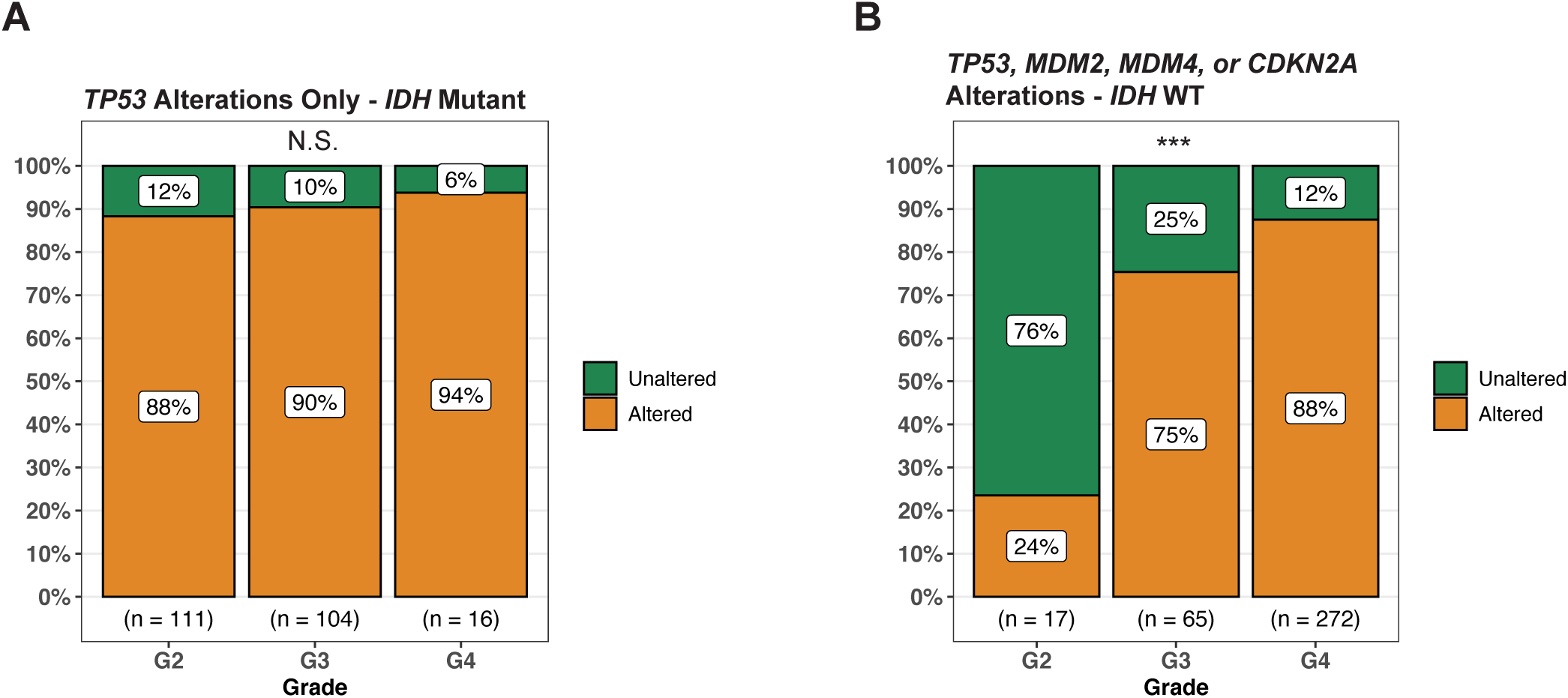
*IDH* WT Glioma progression from low to high grade is marked by an increase in the frequency TP53 pathway alterations. Bar plot showing the frequency of *TP53* alterations by grade in *IDH* mutant TCGA glioma samples. Bar plot showing the frequency of *TP53, MDM2, MDM4* and *CDKN2A* alterations by grade in *IDH* WT TCGA glioma samples. (*** *P* < 0.001; N.S.: not significant, *P* > 0.1).

